# Intracellular genomic variability driven by cellular compartmentalization in the giant bacterium *Achromatium* spp

**DOI:** 10.64898/2026.08.16.745057

**Authors:** Danny Ionescu, Joana Mariz, Kai Bukoff, Elene Tskitishvili, Marit Grüner, Laura Oroz Villegas, Sophie-Luise Heidig, Thomas Stach, Julian Hennies, Dietrich Walsh, Anna M. Steyer, Christian Wurzbacher, Mina Bizic

## Abstract

Bacteria of the genus *Achromatium* harbor hundreds of chromosomes that were previously suggested to be genetically diverse. By sequencing multiple regions from individual cells, we demonstrate that chromosomes within a single cell differ in nucleotide and amino acid sequence, reaching levels of divergence well below accepted bacterial species boundaries and below the range associated with homologous recombination. Sequencing of dividing cells further revealed that daughter cells inherit distinct chromosome populations, a mode of inheritance previously associated with sexual reproduction in eukaryotes. Combining subcellular sequencing with high-resolution microscopy, we reconstruct the three-dimensional cellular architecture of *Achromatium* and show that extensive intracellular heterogeneity arises from the spatial segregation of chromosome populations by the cell’s internal structure, which limits genome-wide recombination. Our findings establish cellular architecture as a determinant of genome evolution in giant polyploid bacteria and identify spatial genome segregation as a mechanism enabling the maintenance and inheritance of divergent chromosome populations in bacteria.

## Introduction

Polyploidy, the presence of multiple chromosome copies within a single cell, is increasingly recognized as a widespread and evolutionarily successful strategy among prokaryotes. While most model bacteria are monoploid or oligoploid, numerous bacterial and archaeal lineages maintain tens, hundreds, or even hundreds of thousands of genome copies per cell ^1–3^. Polyploidy occurs across diverse phylogenetic groups including Cyanobacteria, haloarchaea, methanogenic archaea, and phylogenetically-diverse giant bacteria, indicating that it has evolved repeatedly and provides substantial adaptive advantages ^3,4^. Proposed benefits include increased resistance to DNA damage, buffering against deleterious mutations, enhanced metabolic flexibility, storage of phosphate in genomic DNA, and the ability to maintain cellular function while exploring evolutionary innovation through genetic diversification of chromosome copies^3–5^.

Among prokaryotes, polyploidy reaches its most extreme manifestations in giant bacteria. Members of the genera *Thiomargarita, Epulopiscium, Beggiatoa*, and *Achromatium* contain hundreds to hundreds of thousands of chromosomes and attain cell sizes several orders of magnitude larger than those of typical bacteria^5–7^. In these organisms, polyploidy is thought to overcome fundamental constraints associated with large cell size. Because diffusion becomes increasingly limiting over large intracellular distances, genomes are distributed throughout the cell volume, creating multiple localized centers of transcription and translation that function as decentralized genetic control units. This organization reduces the need for long-range intracellular transport and allows bacteria to achieve dimensions once considered incompatible with prokaryotic cellular organization.

Recent discoveries have further transformed our understanding of giant bacterial cells. The sulfur bacterium *Candidatus* Thiomargarita magnifica, which can reach lengths exceeding one centimeter, was shown to contain its chromosomes and ribosomes within membrane-bound organelle-like compartments termed pepins^8^. This remarkable cellular architecture effectively distributes genetic information throughout the cell and represents one of the most sophisticated examples of intracellular compartmentalization known in prokaryotes. The discovery demonstrated that giant bacteria can evolve highly specialized solutions to overcome diffusion constraints and challenged long-standing assumptions regarding the structural complexity attainable by bacterial cells. However, despite its extraordinary organization, the genome copies of *Ca*. T. magnifica appear largely homogeneous, consistent with extensive coordination and maintenance across the intracellular chromosome population^8^. Whether spatial organization can instead facilitate the persistence of genetic diversity within individual bacterial cells remains unknown.

The evolutionary consequences of maintaining large numbers of chromosomes remain incompletely understood. Classical evolutionary theory predicts that asexual polyploid organisms should experience progressive accumulation of deleterious mutations through Muller’s ratchet, eventually leading to genomic degeneration^9,10^. Yet polyploid bacteria and archaea have clearly escaped this fate. Studies of haloarchaea and other polyploid prokaryotes demonstrated that homologous recombination and gene conversion efficiently eliminate deleterious mutations by replacing defective alleles with functional variants from other chromosome copies^3,4,11^. Gene conversion therefore acts as a powerful homogenizing force, maintaining chromosome integrity while preventing mutational meltdown. In several archaeal species, chromosome copies remain nearly identical despite extensive polyploidy, indicating frequent genetic exchange among coexisting genomes^3,11^.

This prevalence of chromosome homogenization creates an evolutionary paradox. Polyploidy has long been proposed to facilitate evolutionary innovation by allowing the emergence of novel alleles while preserving cellular functionality through redundant chromosome copies. Yet the same mechanisms that protect genome integrity also erase variation through continual gene conversion. Consequently, most characterized polyploid prokaryotes contain chromosome populations that are remarkably uniform within individual cells^3^. Whether extensive and persistent intracellular genomic diversity can exist within a single bacterial cell, and under which cellular constraints such diversity can be maintained, remains largely unknown.

Achromatium represents a striking exception to this pattern. Members of this globally distributed genus of giant sulfur bacteria inhabit freshwater, brackish, and marine sediments and can reach dimensions exceeding 130 × 35 μm^12–15^. Their most conspicuous feature is the presence of large calcium carbonate inclusions occupying the majority of the cell volume. For more than a century, the biological significance of these inclusions remained enigmatic. Recent investigations revealed that they are associated with a highly unusual cellular organization and may play a central role in shaping the physiology and evolution of these organisms ^12,14,16,17^. However, their structural contribution to maintaining this architecture remains unresolved.

Unlike other giant bacteria investigated to date, Achromatium exhibits extraordinary intracellular genetic diversity. Community-resolved and single-cell genomic analyses demonstrated that individual cells harbor large numbers of divergent chromosome copies encoding distinct allelic variants of core genes^12,17^. Subsequent analyses suggested that this diversity extends throughout substantial portions of the genome and reflects the coexistence of genetically differentiated chromosome populations within a single cell. These findings challenged the prevailing view that gene conversion inevitably homogenizes chromosomes in polyploid prokaryotes and established Achromatium as the first documented example of stable large-scale intracellular heterozygosity. Furthermore, comparative analyses of giant bacterial genomes suggested that heterozygosity may not be unique to Achromatium, but could represent a more widespread and previously overlooked feature of giant bacteria^18^.

A potential explanation for this phenomenon emerged from observations of Achromatium cell architecture. Electron microscopy and fluorescence imaging suggested that the large calcium carbonate inclusions occupy a periplasmic compartment, while the cytoplasm is confined to thin interconnected layers surrounding and separating the inclusions^12,16^. Rather than forming a continuous intracellular environment, the cytoplasm may therefore be partitioned into spatially restricted domains. Such an organization could fundamentally alter chromosome dynamics. Gene conversion may occur efficiently among neighboring chromosomes while remaining infrequent between chromosomes separated by large inclusion-filled regions. Under these conditions, localized chromosome populations could evolve semi-independently while retaining sufficient connectivity to maintain overall cellular function.

The implications of such a model extend beyond the biology of Achromatium. If chromosome populations are physically structured within the cell, evolutionary processes normally considered at the level of populations may operate within a single organism. Individual Achromatium cells could behave as spatially organized assemblages of coexisting genomes that experience differential diversification and selection. In this scenario, the bacterial cell becomes an evolutionary arena in which genetic variation is generated, maintained, and sorted below the level of the individual organism. Moreover, because chromosome segregation during cell division would involve partitioning and reshuffling of localized chromosome populations, daughter cells may inherit distinct subsets of intracellular diversity. Such a mechanism would generate genetically unique offspring without requiring mutation or horizontal gene transfer and would represent a previously unrecognized mode of microbial diversification.

In contrast to *Candidatus* T. magnifica, where compartmentalization appears to preserve the coordinated function of a largely clonal chromosome population, intracellular structuring in *Achromatium* may promote the persistence of genomic diversity. These observations suggest that giant bacteria have evolved fundamentally different solutions to the challenges imposed by extreme size, genome maintenance, and intracellular organization. Understanding the relationship between cellular architecture and chromosome dynamics in *Achromatium* therefore provides an opportunity to examine how evolutionary processes can operate within the boundaries of a single bacterial cell.

Here we combine super-resolution fluorescence microscopy, focused ion beam scanning electron microscopy (FIB–SEM), subcellular and single-cell sequencing, and metabolic labeling approaches to investigate the relationship between cellular architecture and genome organization in Achromatium. By sequencing physically separated regions of individual cells and naturally dividing cells, we directly assess the spatial distribution and inheritance of genetic diversity. In parallel, incorporation of EdU, EU, and BONCAT enables visualization of DNA synthesis, RNA synthesis, and protein synthesis at subcellular resolution. Together, these approaches reveal that the extraordinary heterozygosity of Achromatium is intimately linked to its highly structured cellular architecture, providing a mechanistic explanation for how extensive intracellular genomic diversity can be maintained within a single bacterial cell. Our findings establish Achromatium as a model system for studying genome evolution below the cellular level and broaden current concepts of individuality, inheritance, and adaptation in giant polyploid bacteria.

## Results

### Sub-cellular chromosomes dissimilarity

To investigate the intracellular genomic diversity, multiple sections from 12 *Achromatium* cells were extracted using laser dissection microscopy, sequenced, and their average nucleotide (ANI) and amino acid (AAI) identity was compared. Overall, between 2 and 9 cell parts were successfully amplified. The assembly of these reads, following removal of non-Achromatium sequences (see methods) ranged between 36 KB and 13,580 KB with an average of 1,530 KB and a median of 301 KB (Table S1). Similarly, 8 sister cells were split in mid division, sequenced separately and compared. These assemblies ranged between 31 KB and 9,290 KB with an average of 3,307 KB and a median of 2,784 KB.

ANI between cell parts was often well below the values accepted for genomes within a single bacterial species (i.e. <94 %). ANI between different parts of the same cell ranged between 81.7 % and 99 % when only alignments above 100 KB were considered (Fig. 1A) but reached also 79% when the minimum alignment length was reduced to 20 KB. In some cells (e.g. cell 5; Fig. 1B) discrete clusters of highly similar (ANI ∼99 %) were observed. These pairs are strongly divergent, with an inter-pair ANI of ca. 91 %. In other cells, e.g. cell 3, almost no variability in ANI values was observed between cell parts, being in all cases above 99 % similar, though no clonal chromosomes were observed in any of the cells.

**Figure. 1.**
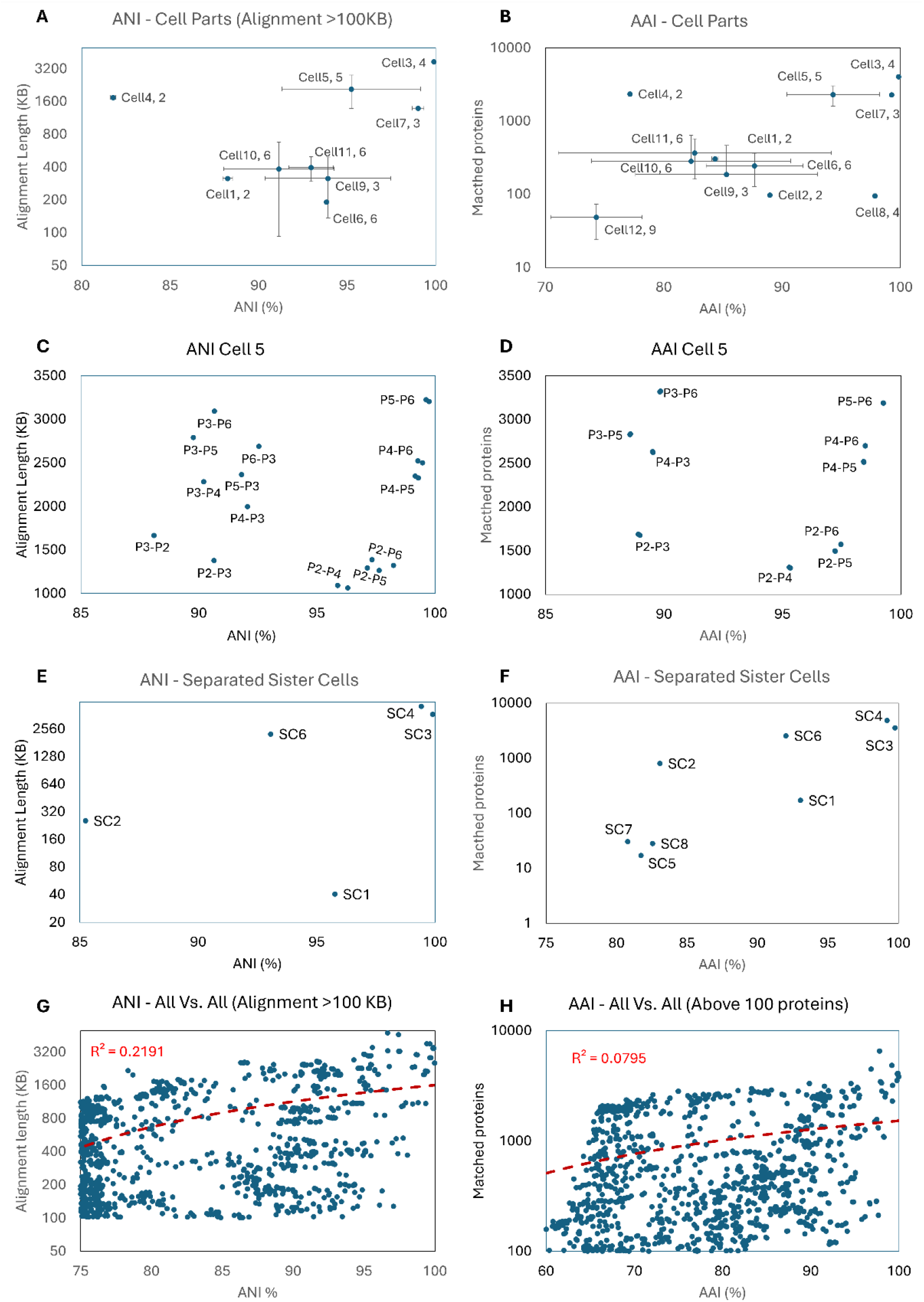
An overview of Average Nucleotide (ANI, A) and Average Amino-Acid (AAI, B) Identity between parts of *Achromatium* cells; a focus on a single cell (C, D). ANI and AAI between sister cells separated mid division are shown in panels E and F, respectively. The dependence of ANI and AAI values on alignment length or the number of homologous proteins is shown in panels G, and H, respectively. In panels A and B the labeling refers to the cell number and the number of cell parts successfully amplified (e.g. Cell11,6 means 6 parts from Cell11 were sequenced). In panel C the labeling indicates the part number compared (e.g. part5 vs. part6 are labeled P5-P6)

Split sister cells exhibited a similar variability in ANI ranging between 85 % and 99 % (Fig. 1C). Analysis of all ANI comparisons (Fig 1H) reveals only weak correlation between ANI and alignment length confirming validity of the observed low intracellular ANI values. Average Amino Acid (AAI) analysis, mirrored the ANI results revealing similar trends of high intracellular diversity (Fig. 1D-F).

Using and all versus all comparison within our dataset, we evaluated the effect of alignment length or number of identified homologous proteins on ANI (Fig1G), or AAI (Fig1H), respectively. The analysis revealed a weak (ANI) to non-existing (AAI) correlation between the amount of data available for comparison and similarity measure, within the range used.

### Physical separation between cellular compartments

Three-dimensional segmentation models conducted using super-resolution fluorescent (Fig. 2A) and Cryo Focused Ion Beam Scanning Electron Microscopy (FIB-SEM; Fig. 2B) images, reveal a complex internal structure consisting of small compartments, almost entirely enclosed by the cytoplasmatic membrane. The cell’s chromosomes lie in these compartments (Fig. 2A), as could be shown through the colocalization of >90 % of the DNA spots with membranes.

**Fig. 2.**
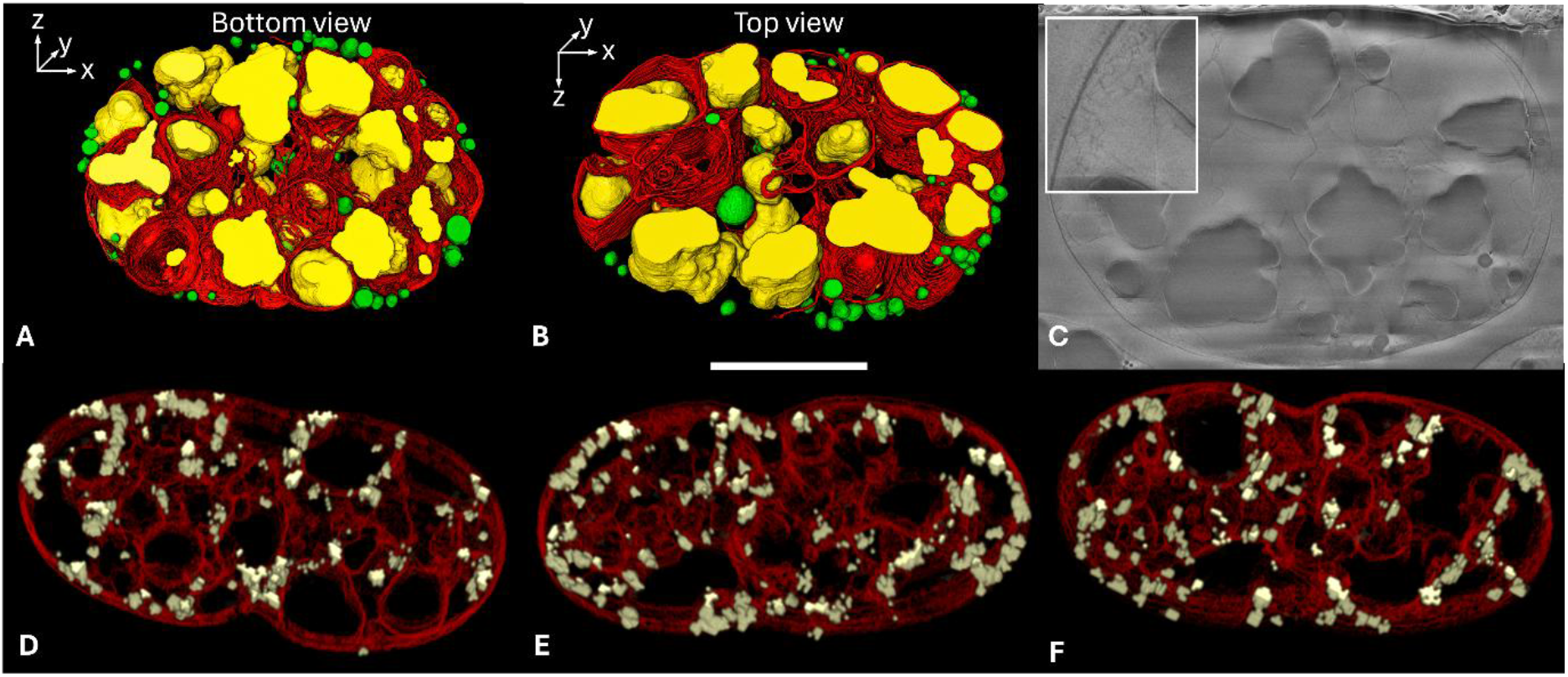
Detailed cellular architecture of *Achromatium* cells depicting the complex membranal structure and intracellular compartmentalization. Panels (A) and (B) are side and top views, respectively, of a 3D segmentation model generated from half a cell imaged with cryo-FIB-SEM with a voxel size of 5×5×30 nm (x,y,z). Membranes are labeled in red, CaCO_3_ in yellow and sulfur globules in green. In total ca. 450 slices were used for the model. An example of a single slice is shown in panel (C) The cell periphery is characterized by an elaborate network of membranes that were highlighted and magnified in the insert in panel (C) and was not segmented. Outer membrane and cell wall are not featured in the segmentation model. Panels D-E depict 2.5 μm thick slabs segmented from super-resolution imaging of DNA (yellow) and membranes (red) of a dividing *Achromatium* cell. The localization of the DNA in spatially separate membranal pockets is evident along all three (x,y,z) axes. Panels A-C and D-F are derived from difference specimens. The white scale bar is 20 μm long.

DNA staining of multiple discrete DNA signals distributed throughout the cell confirmed previous results^12,16^, representing individual chromosomes and chromosome clusters (Figure 2D-E, 3A). Quantitative analysis identified between 107 and 353 distinct DNA foci per cell. The number of chromosomes was positively correlated with cell size and surface area. The average chromosome density was one chromosome per 71±29 μm^3^. Each DNA focus occupies, on average, 20±8 μm^3^ of CaCO_3_-excluded cellular volume. However, the median inter-focus distances is 2.1±1 μm, resulting in a local chromosome-centered interaction domain of 3–6 μm^3^.

A pronounced diversity in morphology among *Achromatium* cells was observed, varying in cell size, shape, and in the morphology of their calcite bodies. Cells ranged from 32 μm to 95 μm in length. SEM analyses further highlight heterogeneity in the size of intracellular calcite inclusions, which range from large, well-defined bodies (up to 10 μm in diameter) to smaller, more granular structures (down to 1 μm in diameter). Interestingly, FIB-SEM reveals that membranes do not necessarily closely follow the calcite bodies, suggesting that changes in calcite body size does not immediately result into resizing of membranal compartments.

To assess the contribution of the CaCO_3_ bodies to intracellular architecture, Nile Red-stained cells were followed during HCl-induced dissolution of the CaCO_3_ inclusions (Fig. S1). Membrane-associated fluorescence persisted following decalcification; however, the characteristic cellular shape and organization of the membrane-stained structures were progressively lost. Thus, removal of the CaCO_3_ bodies was associated with disruption of the intracellular architecture without loss of membrane staining.

### Localization of cellular processes

Given the observed compartmentalization we asked whether basic cellular processes such as DNA synthesis (replication), transcription, and translation, occur uniformly across the cell or are these processes restricted to a few compartments.

DNA synthesis occurs throughout the cell, yet *Achromatium* does not replicate all its chromosomes in each replication cycle (Fig. 3A-C). EdU incorporation during DNA replication revealed several discrete activity sites across the cell, always fewer in number compared to the general DNA staining signals, with an average of 58±21 % (n_cells_=25) of chromosomes being replicated. In contrast transcriptional and translational activity occurs throughout the cell. Newly synthesized RNA (Fig 3D-F) and proteins (Fig. 3G-I) showed diffuse patterns, with several foci in close proximity to chromosomes in the case of RNA synthesis. No similarity was observed between localized DNA replication patterns and transcriptional and translational activity which occurred throughout the entire cell.

**Figure 3.**
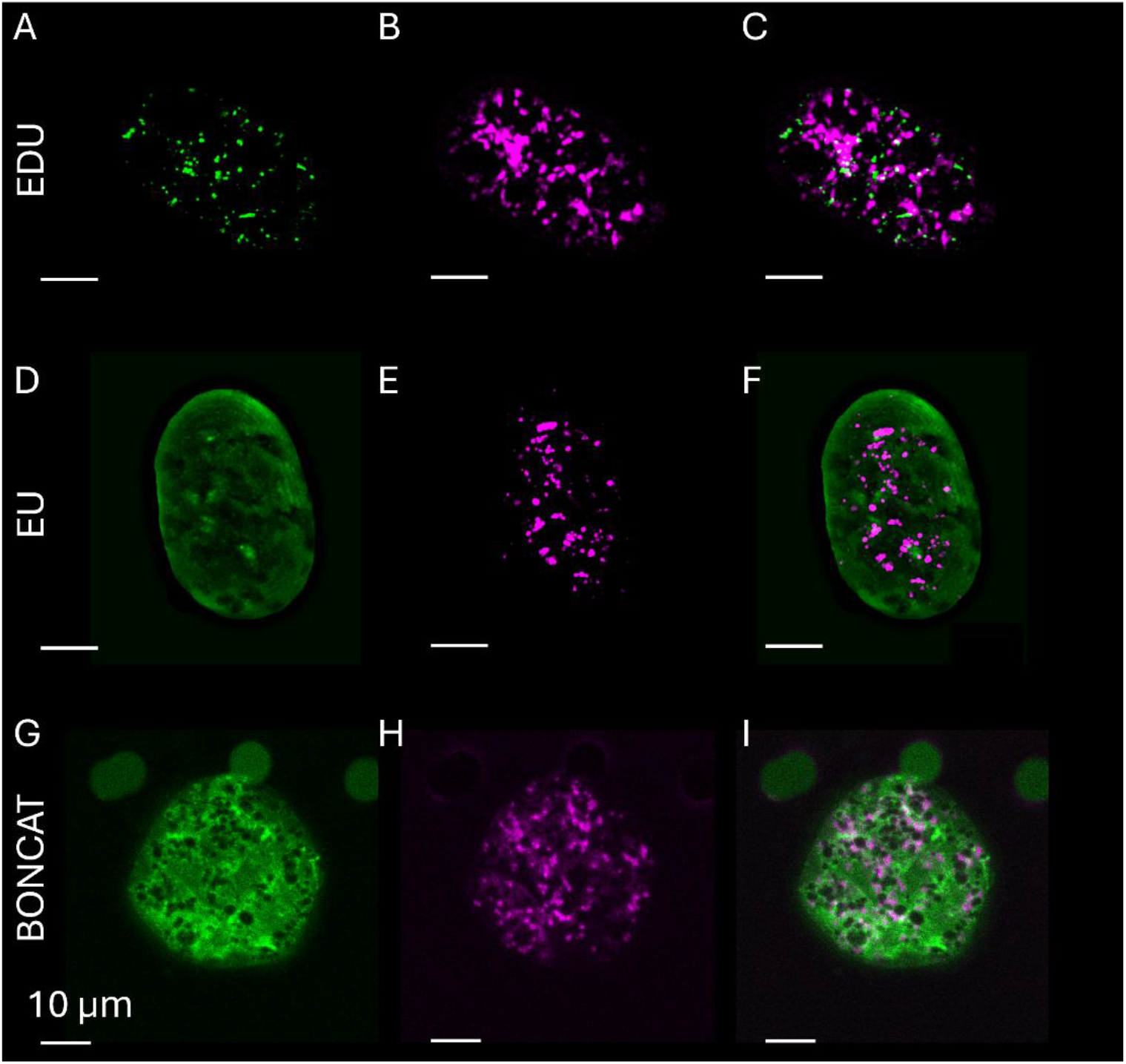
Confocal laser scanning microscopy of cell labeled with Edu for DNA synthesis detection (A-C), Eu for RNA synthesis detection (D-F) and BONCAT for protein synthesis (G-I). Green channels show the specific assays, and magenta channels show total chromosomes staining using DRAQ5. Panels C,F,I show overlaid images for each assay. Labelling controls with killed cells for newly synthesized DNA, RNA and proteins exhibited no detectable EdU, Eu, or BONCAT signals, respectively.

## Discussion

Our results provide direct evidence that the remarkable intracellular genomic diversity previously described in *Achromatium* arises from, and is maintained by, its unique cellular architecture. By combining subcellular sequencing with three-dimensional ultrastructural imaging and metabolic labeling, we demonstrate that chromosomes are physically partitioned into membrane-delimited cytoplasmic pockets, creating spatially isolated chromosome populations that can diverge extensively within a single bacterial cell. The observation that different regions of the same cell frequently exhibit average nucleotide identities below accepted bacterial species boundaries, while daughter cells inherit distinct subsets of this diversity, fundamentally changes our understanding of genome organization in giant polyploid bacteria.

### Cellular compartmentalization prevents global genome homogenization

Polyploid bacteria generally maintain highly homogeneous chromosome populations despite carrying tens to hundreds of genome copies. This homogeneity results from continual homologous recombination and gene conversion that prevent the accumulation of deleterious mutations predicted by Muller’s ratchet^3,4,11^. Our observations suggest that *Achromatium* has largely escaped this homogenizing force.

The genomes within *Achromatium* appear to form multiple semi-independent evolutionary units whose interactions are constrained by cellular architecture. Achromatium’s cytoplasm was shown to occupy the spaces in between the CaCO_3_ bodies^16^. Here, we reconstructed the 3D structure of the cell using Cryo-FIB-SEM and super-resolution microscopy and reveal that the cytoplasm is subdivided into numerous membrane-defined compartments. DNA foci (i.e. one or more chromosomes) colocalize with these disconnected membrane pockets and appear to have no physical connectivity with each other beyond their immediate subcellular surroundings (see. Fig. 2 D-E, and supplementary movie). The CaCO_3_ bodies appear to contribute to maintaining this organization: following their experimental dissolution, membrane-associated fluorescence persisted, but the characteristic intracellular architecture was disturbed (Fig. S1). This supports a structural role for the CaCO_3_ bodies in maintaining the spatial organization of the membrane-defined cytoplasmic compartments.. Under these conditions, homologous recombination is expected to occur predominantly among chromosomes within the same local compartment, whereas exchange between distant chromosome populations is eliminated or greatly reduced. The previously suggested activity of mobile genetic elements^12^ likely operates within these compartments giving rise to the inconsistent gene synteny observed in *Achromatium* genomes^12^. Although our data cannot exclude occasional long-range genetic exchange, e.g. in the rare cases when the CaCO_3_ are temporarily absent^16^, they strongly suggest that chromosome homogenization operates locally rather than across the entire cell. Overall, this explains why extensive heterozygosity can persist despite extreme polyploidy. Rather than behaving as a single chromosome population,

### Cellular architecture generates evolutionary structure below the level of the organism

Subcellular sequencing revealed remarkable genetic differentiation between physically separated chromosome populations. In several cells, chromosome amplified from different parts of the cell exhibited near-clonal similarity while differing substantially from those from other parts. This suggests prolonged local evolution, however at non-uniform rates across the cell, likely driven by selective chromosome replication. Similar levels of divergence were also detected between daughter cells separated during cytokinesis, demonstrating that cell division partitions this diversity asymmetrically.

These observations support a model in which chromosome populations evolve locally during vegetative growth but are partially reshuffled during cell division. As septation progresses, previously separated chromosome compartments become transiently connected, allowing limited mixing before daughter cells inherit new combinations of chromosome clusters. Such a mechanism resembles, conceptually, a form of genetic reassortment rather than classical bacterial inheritance. Although distinct from meiotic recombination, this process generates genetically unique offspring without requiring new mutations or horizontal gene transfer.

This represents a previously unrecognized mode of diversification in bacteria, where evolutionary processes normally associated with populations occur within individual cells. The bacterial cell therefore becomes not only the unit of selection but also an arena in which standing genomic variation is continuously generated, maintained, and reorganized.

### Selective replication and functional heterogeneity

Our metabolic labeling experiments further suggest that chromosome populations are regulated independently. Only approximately 60% of chromosome foci incorporated EdU during the experimental period, indicating that chromosome replication is selective rather than synchronous. In contrast, transcription and protein synthesis occurred throughout the cell, indicating that all cellular compartments are active, regardless of DNA replication activity. Similar asynchronous replication has been observed in other polyploid bacteria and archaea^3,19^, but the combination of selective replication with extensive intracellular heterogeneity appears unique to *Achromatium*.

Selective DNA replication could serve as a long-term adaptation strategy to new environments or to ongoing constant changes. It was previously hypothesized that the intracellular allelic diversity offers *Achromatium* a mechanism to fine tune it’s response to environmental conditions^12,17^, a feature previously shown for eukaryotes^20–22^. A large survey for Achromatium genes expressed under different environmental conditions conducted with public metatranscriptomes revealed differential gene expression between freshwater and marine Achromatium, despite lack of characteristic functional enrichment^17^. A parallel metaproteomics study on *Achromatium* reveals temperature dependent expression of protein homologs supporting this hypothesis (**Heidig et al**., **submitted**). We further hypothesized that long term adaptation could be done through gene copy enrichment of specific alleles^17^, similar to eukaryotes^23–25^. Given the large cells size and the centrally localized cell division site, DNA replication in *Achromatium* is not spatially or temporally linked to cell division. The latter is evident from DNA replication in non-dividing cells, and away from the cell’s center. The ability to selectively replicate some chromosomes over others, regardless of their distribution between daughter cells would allow *Achromatium* to enrich beneficial alleles, while maintaining alternative ones as archived backups for alternating environments or conditions. It is important to note that at this point, we do not have evidence linking between selective chromosome enrichment, specific alleles, and environmental change.

### Comparison with other giant bacteria

Compartmentalization has recently emerged as a defining feature of several giant bacteria. In *Candidatus* Thiomargarita magnifica, chromosomes and ribosomes are enclosed within membrane-bound structures termed pepins, effectively creating distributed centers of transcription and translation throughout the enormous cell^8^. In contrast, available evidence suggests that chromosome copies in *Ca*. T. magnifica remain highly homogeneous. A similar chromosomal homogeneity was shown for the polyploid filamentous bacterium *Candidatus* Marithrix sp.^26^. In contrast the unicellular giant bacterium *Epulopiscium* sp. was suggested to host heterogenous chromosomes as well^18,27^. The difference between levels of chromosome homogeneity in different polyploid giant bacteria is likely attributed to the means of cells replication ^18^. Binary fission or spore forming in unicellular giant bacteria promotes physical segregation between chromosomes from different parts of the cell. In contrast filaments elongate in one or two directions from a single point, generating clonal chromosome copies in parallel to their growth.

*Achromatium* therefore represents a fundamentally different solution to the challenges imposed by extreme cell size. Rather than merely overcoming diffusion limitations, compartmentalization appears to generate spatial isolation sufficient to promote long-term genomic diversification. These contrasting strategies suggest that giant bacteria have independently evolved different organizational principles balancing chromosome maintenance, cellular coordination, and evolutionary innovation.

### Implications for bacterial individuality

Our findings challenge traditional concepts of bacterial individuality. A bacterial cell is generally assumed to contain genetically equivalent chromosomes, making the cell itself the fundamental unit of inheritance. In *Achromatium*, however, chromosomes within the same cytoplasm may differ by genetic distances exceeding accepted species boundaries. Yet these diverse genomes cooperate to maintain a single physiologically integrated organism.

This organization resembles a structured population more than a conventional bacterial cell. Cellular architecture establishes barriers to genetic exchange analogous to geographic isolation in multicellular populations, allowing chromosome populations to diversify while remaining functionally integrated. Similar principles operate in population genetics at much larger spatial scales; *Achromatium* compresses these evolutionary dynamics into the interior of a single prokaryotic cell.

More broadly, these observations suggest that the relationship between cellular organization and genome evolution deserves greater consideration across polyploid microorganisms. If intracellular compartmentalization can influence evolutionary trajectories, then selection, diversification, and inheritance may operate simultaneously at multiple organizational levels within a single organism.

Future work combining live-cell imaging, chromosome tracking, and functional genomics will be required to determine how chromosome populations are maintained, how frequently they exchange genetic material, and whether distinct chromosome clusters occupy stable physiological niches. Understanding these processes may ultimately redefine how individuality, inheritance, and adaptation are viewed in giant bacteria and perhaps in polyploid prokaryotes more generally.

## Methods

### Cell collection

*Achromatium* cells were collected from Lake Stechlin in Brandenburg, Germany (53°08’29.7”N, 13°01’46.8”E). Sampling took place in mid-May, late June, and at the end of August. Surface sediment was collected using a mug and transferred to jars, filling them to a depth of approximately 4 cm with sediment, the remaining volume was filled with lake water, leaving an air gap to maintain oxygen availability. They were stored with a loosely closed lid to allow gas exchange, either at room temperature or at 6°C in the refrigerator. For staining procedures, *Achromatium* cells were freshly harvested from sediment samples kept in jars. To isolate cells and remove large debris, the upper sediment layer was pipetted and sequentially filtered through a series of sieves with decreasing pore sizes (200 μm, 125 μm, 100 μm, and 63 μm). The filtrate was collected in a glass Petri dish for further processing. Individual cells were then isolated under a stereomicroscope by rotating and tilting the Petri dish against a dark background with lateral illumination. *Achromatium* cells were subsequently transferred to a clean black glass microscopy dish.

### Sequencing

Achromatium cells were laser-dissected using an LMD7 for single-cell dissection, as previously described^28^. To analyze dividing Achromatium cells, the cells were washed and distributed on a UV-irradiated PPS membrane (30 minutes at 11.7 J/cm^2^) coated with poly-L-lysine (0.01%) and briefly dried at 27 °C for 15 minutes. The membrane slide was pre-screened to identify dividing cells based on their morphology. The laser was then used to separate the cells into two parts, which were subsequently subjected to whole genome amplification and Illumina sequencing (NextSeq2000), without the zymolase step, as described in Mariz et al., 2025. To analyze Achromatium cell parts, we distributed the cells on the membrane as described above. The full slide was then screened for isolated single cells and marked their positions. The slide was then placed in a sandwich setup within a clean bench, with a UV-irradiated slide holder at the bottom and a glass slide at the top to break up the Achromatium cells by applying pressure. The membrane slide was then transferred back to the LMD, where the cell parts of a single Achromatium cell were dissected for WGA, as described above.

### Bioinformatic analysis

Raw reads from *Achromaium* single cells or cell parts were quality trimmed using Trimmomatic^29^ (V.) and subsequently assembled using Spades^30^ (V) using the single cell flag. To obtain assembiles consisting only of *Achromatium* data, the following approach was taken. First, binning of the assemblies was performed using Metabat2^31^. Bins identified as *Achromatium* using GTDB-tk^32^ (DB v. 226) were kept and were merged per sample. In parallel, all contigs were taxonomically annotated using the CAT-Pack tool^33^ using the same GTDB reference database. Un-binned contigs matching *Chromatiales, Chromatiaceae*, and *Achromatium* were kept, while binned contigs, specifically associated with other known taxa (at any taxonomic depth) were removed. Finally, the contigs considered to belong to *Achromatium* were consolidated into one bin per sample. Average nucleotide identity was conducted using the FastANI^34^ (v. 1.34) tool comparing all obtained *Achromatium* mags one to another. Average Amino-acid Identity was calculated using EZAAI^35^ (v. 1.2.4) tool following protein calling using Prodigal^36^ (v. 2.6.3).

### Super Resolution Microscopy

Super Resolution Images were acquired on a Leica Stellaris 8 STED FALCON system equipped with a first generation white light laser, a 775nm pulsed STED laser, and the FALCON fluorescence lifetime imaging module.

Samples were prepared and allowed to settle to the bottom of Ibidi chambers (35mm and 8well) with 1.5H high precision glass bottoms before being imaged using confocal and STED modalities.

Excitation, reflection and STED laser intensities were optimized on a sample by sample basis to achieve the best possible results. All specific settings (Laser wavelengths and intensities, detector width, pinhole, scan speed, accumulations) on an image by image basis are in the metadata of the raw files which are available on request.

### Activity staining

For incubation with the corresponding labeling reagent, the collected cells were placed onto a filter (pore size 4 μm) and positioned on top of a second filter (pore size 2 μm). The filter stack was placed on sediment inside a vacuum filtration unit. The sediment surface was sealed with aluminum foil to prevent solutions from flowing through during incubation. After incubation, the foil and sediment were removed, and the filtration unit was connected to a vacuum pump. All subsequent solutions were drawn through the filters using vacuum filtration.

### EDU Staining

To stain newly synthesized DNA, the Click-iT™ EdU Alexa Fluor™ 488 Imaging-Kit (Invitrogen) was used according to the manufacturer instructions. Shortly, a 10 mM EdU stock solution was diluted in filtered lake water to a final concentration of 10 μM. Cells were incubated with EdU at room temperature or at 6°C in the refrigerator for 24 hours. After incubation, the sediment and the remaining solution were removed. Cells were fixed in 2% formaldehyde (Polysciences, Inc.) in 1X phosphate-buffered saline (PBS, 10X: 80 g/l *NaCl*, 2 g/l *KCl*, 17.8 g/l *Na*_2_*HPO*_4_ · 2*H*_2_*O*, 2.4 g/l *KH*_2_*PO*_4_) for one hour at room temperature. The fixative was removed, and the cells were washed twice with 1 ml of 3 % bovine serum albumin (BSA) in PBS, after which they were permeabilized with 200 μl of 0.5 % Triton™ X-100 (Sigma-Aldrich) in PBS for 30 minutes at room temperature. Following permeabilization, cells were washed twice with 3 % BSA in PBS. To minimize non-specific binding, cells were incubated with 1 ml of ROTIBlock (Carl Roth), diluted 1:10 in ultrapure H_2_O, for 1 hour while gentle shaking. After removing the blocking solution, the reaction cocktail was prepared according to the kit protocol and added to the cells, adjusting the final volume to 200 μl while maintaining the recommended component ratios. Cells were incubated with the reaction mixture for 30 minutes at room temperature in the dark, followed by a final wash with 1 ml PBS. Subsequently, DNA staining was performed.

### EU staining

RNA was stained using the Click-iT™ RNA Alexa Fluor 488™ Imaging-Kit (Invitrogen). A 100 μM EU working solution was prepared by diluting in filtered lake water. Cells were incubated with the reagent for 20 hours at either room temperature or 6 °C in the refrigerator. Subsequent fixation, permeabilization, and blocking procedures followed the same protocol as described above for DNA labelling. Cells were washed with PBS instead of BSA. The reaction cocktail was prepared according to the kit protocol, using a total volume of 200 μl while maintaining the prescribed reagent ratios. After incubation for 30 minutes at room temperature in the dark, cells were washed with 1 ml of the provided reaction rinse buffer. Subsequently, DNA staining was performed.

### BONCAT

For protein synthesis labeling, bioorthogonal non-canonical amino acid tagging (BONCAT) was performed using the Click-iT™ AHA Alexa Fluor™ 488 Protein Synthesis HCS Assay (Invitrogen). Cells were incubated with 50 μM AHA diluted in filtered lake water for 20 hours at either room temperature or 6 °C. Following incubation, AHA was removed, and cells were washed with 1 ml PBS. Cell fixation, permeabilization, and blocking were performed as described above for DNA synthesis labelling. The reaction cocktail was prepared according to the manufacturer’s protocol, using a total volume of 200 μL while maintaining the original reagent ratios. After a 30-minute incubation at room temperature in the dark, cells were washed with 1 ml 3 % BSA in PBS and then processed for DNA staining.

### Total DNA Staining for metabolic activity imaging

DNA was stained using Deep Red Anthraquinone 5 (DRAQ5) (Miltenyi Biotec). Cells were resuspended in a 15 μM DRAQ5 solution, diluted in PBS and incubated for 30 minutes at room temperature in the dark. Cells were washed once with PBS.

The samples were imaged immediately using a Leica TCS SP5 confocal laser scanning microscope (CLSM) or stored at –20 °C and imaged within one week.

### *Escherichia coli* positive control

*Escherichia coli* W1485 was used as a positive control. Cells were grown in LB medium (EdU and EU experiments) or methionine-free M9 minimal medium (AHA experiments) at 37 °C to mid-exponential phase (OD_600_ = 0.4–0.6). Cultures were incubated with 10 μM EdU for 15 min, 100 μM EU for 15 min, or 100 μM L-azidohomoalanine (AHA) for 30 min at 37 °C with shaking. Cells were pelleted by centrifugation (11,000 rpm, 2 min) between labeling and washing steps, replacing the vacuum filtration used for *Achromatium*. Subsequent click-chemistry labeling, membrane staining, and fluorescence imaging were performed as described for *Achromatium*. DNA was counterstained with DAPI (1 μg ml^−1^, 5 min). Cells were mounted on 0.1% gelatin-coated slides using antifade mounting medium and imaged by fluorescence microscopy. For negative controls, cells were fixed prior to addition of the respective labeling reagent, with all subsequent steps performed identically.

## Image analysis

### EDU labelling analysis

Raw Z-stacks from channels 1 (525 nm emission) and 3 (700 nm emission) were processed independently. Point spread functions were generated using PSF Generator (Kirschner et al., 2013) with Born & Wolf 3D optical model (NA 1.3, pixel size 286 nm, Z-step 339 nm), and images were deconvolved using DeconvolutionLab2 (Sage et al., 2017). Background was subtracted using a rolling ball algorithm and images converted to 16-bit. Cells were manually selected via ROI manager, thresholded using the Yen algorithm, and segmented using watershed to separate chromosomes. Individual chromosomes were quantified using 3D Object Counter (ImageJ/Fiji), with manual removal of artifacts using the wand tool when necessary.

Images of Achromatium cells from DNA, RNA and Protein synthesis assays were captured as z-stacks to enable 3D reconstruction. The resulting image stacks were processed and analyzed using Fiji (ImageJ distribution, fiji), including basic contrast adjustments and quantitative 3D analysis.

### Segmentation analyses

#### Amira

A FIB–SEM dataset, comprising 419 consecutive z-sections representing half an *Achromatium* cell, was imported into Amira 6.4.0 (Thermo Fisher Scientific). The voxel dimensions were defined according to the acquisition parameters as 10 × 10 × 30 nm^3^ in the x-, y-, and z-directions. Peripheral regions of the image stack were cropped in order to restrict the dataset to the area of interest. The image volume was then subjected to rigorous inspection in the orthogonal XY, XZ, and YZ planes using the slice-viewing functions of Amira. Manual segmentation was performed slice-by-slice in the Segmentation Editor. The assignment of separate materials within a Label Field to three morphologically distinguishable internal structural components was achieved. The cell envelope was intentionally excluded from the segmentation. Segmentation boundaries were reviewed across consecutive z-sections and in the orthogonal views to maintain spatial continuity and minimize inconsistencies between neighboring slices. The three materials were assigned distinct colours in the material list and subsequently visualised as a combined three-dimensional reconstruction using the Volren volume-rendering module. Material-specific colour and opacity settings were adjusted to display the complex arrangement of the segmented structures.

## Data availability

The sequence data generated in this paper is available on the SRA under Bioproject number PRJNA1538148 at https://www.ncbi.nlm.nih.gov/bioproject/1538148.

## Acknowledgements

The study was funded through DFG grant “**Eco-evolution and cellular functionality of the heterozygous bacterial genus Achromatium**” (project number 465407921) awarded to DI.s

We acknowledge the access and services provided by the Imaging Centre at the European Molecular Biology Laboratory (EMBL IC), generously supported by the Boehringer Ingelheim Foundation.

## Supplementary Material

**Figure S1.**
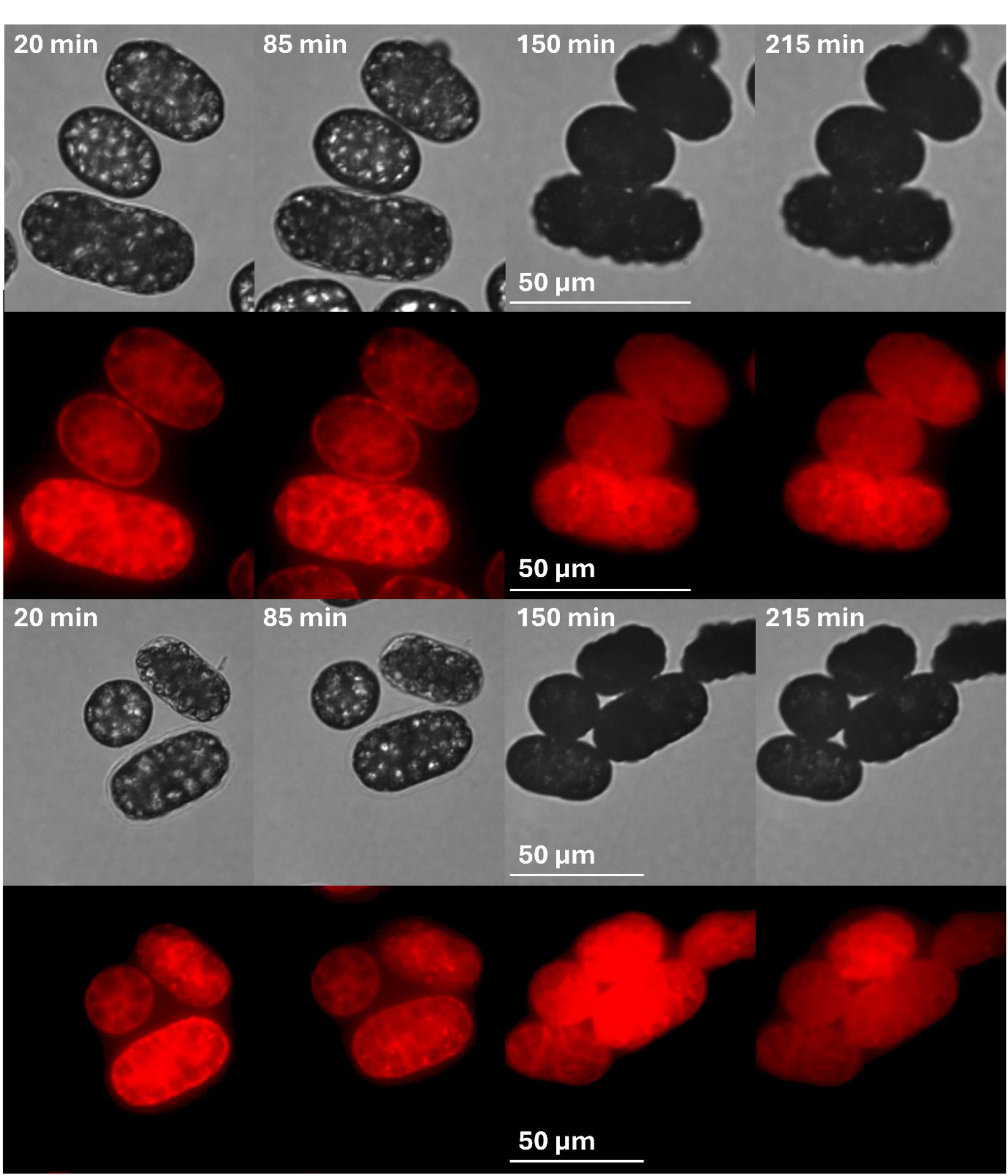
CaCO_3_ dissolution disrupts the intracellular organization of *Achromatium*. Time-lapse microscopy of Nile Red-stained cells during incubation in 5 mM HCl. The upper and lower pairs of image series show two fields of cells from the same experiment, with transmitted-light images above the corresponding Nile Red fluorescence images. Cells moved slightly during incubation and imaging, resulting in changes in their relative positions and the appearance of additional cells within the field of view. Following dissolution of the refractile CaCO_3_ bodies, which are no longer visible in transmitted light, membrane-associated fluorescence persists but the characteristic intracellular organization becomes increasingly diffuse.

