## Supplementary figures and images for "Intracellular genomic variability driven by cellular compartmentalization in the giant bacterium *Achromatium* spp"

### Supplementary Figure 1

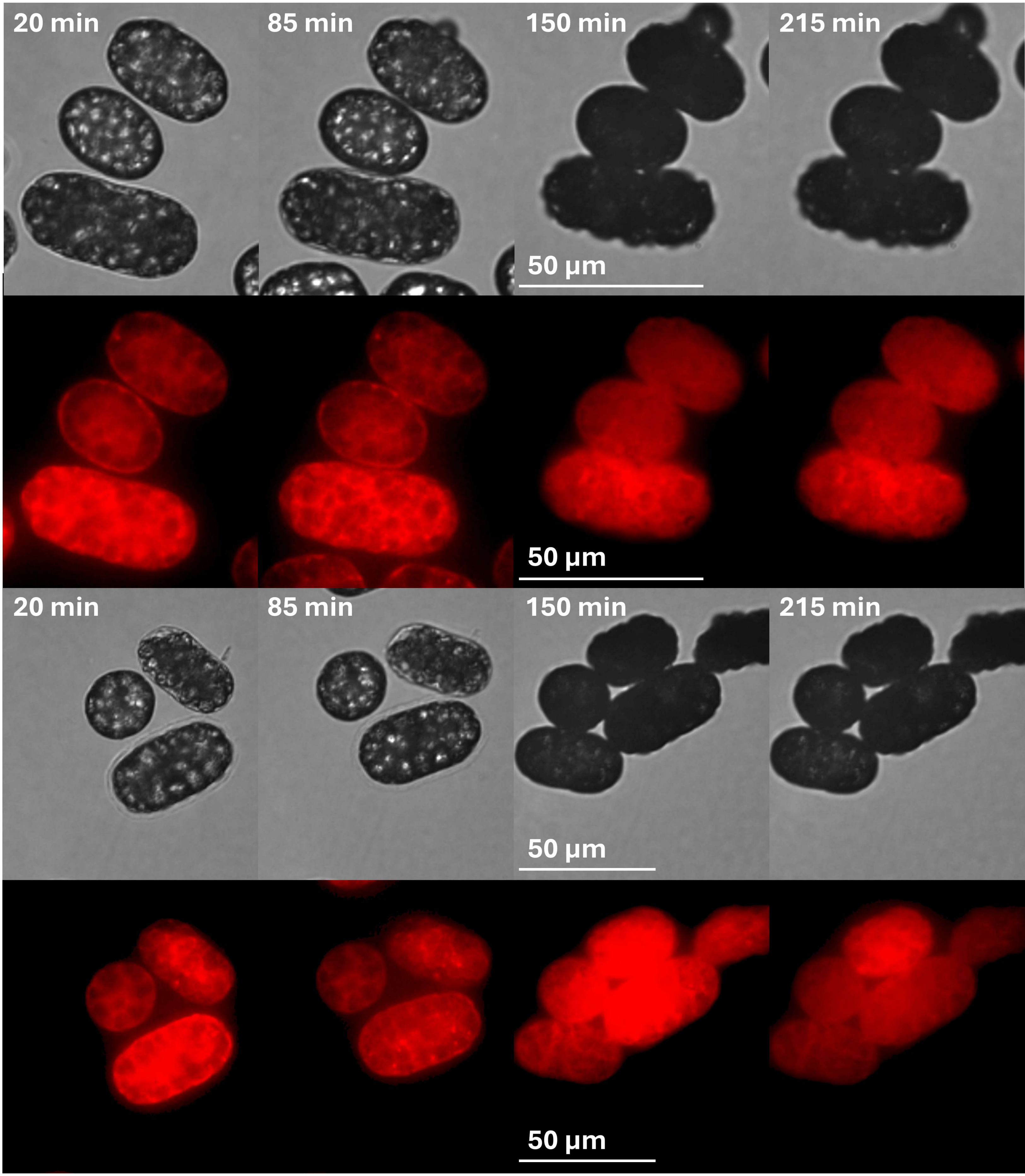
